# Physiological preparation of hair cells from the sacculus of the American bullfrog (*Rana catesbeiana*)

**DOI:** 10.1101/121582

**Authors:** Julien B Azimzadeh, Joshua D Salvi

## Abstract

**SHORT ABSTRACT:** The American bullfrog’s sacculus permits direct examination of hair-cell physiology. Here we describe the dissection and preparation of the bullfrog’s sacculus for biophysical studies. We show representative experiments from these hair cells, including the calculation of a bundle’s force-displacement relation and measurement of its spontaneous motion.

**LONG ABSTRACT:** The study of hearing and balance rests upon insights drawn from biophysical studies of model systems. One such model, the sacculus of the American bullfrog, has become a mainstay of auditory and vestibular research. Studies of this organ have revealed how sensory cells—hair cells—can actively detect signals from the environment. Because of these studies, we now better understand the mechanical gating and localization of a hair cell’s transduction channels, calcium’s role in mechanical adaptation, and the identity of hair-cell currents. This highly accessible organ continues to provide insight into the workings of hair cells.

Here we describe the preparation of the bullfrog’s sacculus for biophysical studies on its hair cells. We include the complete dissection procedure and provide specific protocols for the preparation of the sacculus in specific contexts. We additionally include representative results using this preparation, including the calculation of a hair bundle’s force-displacement relation and measurement of a bundle’s spontaneous motion.

## INTRODUCTION

The acousticolateralis organs of mammals possess a complex architecture and lie within an anatomical niche that can be difficult to access. For example, the mammalian cochlea comprises a spiraling labyrinth and is embedded within the thick temporal bone. Isolation of the cochlea often causes mechanical damage to the sensory cells lying within it and has therefore proven to be a difficult task^1^. Neuroscientists have thus turned to model systems which are more readily extracted from the sanctum of the ear.

One of these model systems, the sacculus of the American bullfrog (*Rana catesbeiana*), has for decades yielded generalizable insight into the function of auditory and vestibular systems. The sensory cells of the sacculus are its hair cells, specialized transducers that convert mechanical energy into electrical signals within our auditory and vestibular organs. Projecting from the apical surface of each hair cell is a mechanosensitive hair bundle that comprises a graded tuft of enlarged microvilli called stereocilia. The tips of adjacent stereocilia are interconnected by filamentous tip-link proteins that mechanically gate ion channels in response to mechanical stimuli^2,3^. Although auditory and vestibular organs respond to different types of stimuli, they share a common detection mechanism. This commonality underlies the many insights gained into hair-cell mechanotransduction through studies of the bullfrog sacculus. For example, the hair cell’s active process has been studied extensively in this organ^4-7^. Not only has it been shown that hair cells generate active work^6^, but distinct mechanisms underlying the active process and a hair cell’s tuning characteristics have been unveiled through studies of bullfrog acousticolateralis organs. These include active motility of hair bundles^8^, electrical resonance within hair-cell somata^9-11^, and frequency selectivity at the hair cell’s ribbon synapse^12^.

The bullfrog’s sacculus appeals to sensory neuroscientists for numerous reasons. Unlike the mammalian cochlea, this organ lies within the readily accessible otic capsule. Second, hair cells within this organ can remain healthy for several hours under appropriate conditions^13,14^. This permits experimentation on these cells over long timescales relative to their mammalian counterparts. Third, the organ bears little curvature, permitting easy manipulation. Fourth, each organ comprises a thousand or more hair cells^15^, providing both a high throughput and a high probability of locating an appropriate set of hair cells for a given experiment. Finally, the bullfrog’s sacculus is easily visualized due to the thinness of this organ and large size of its hair cells.

These properties provide great versatility for the study of sensory cells within the bullfrog’s sacculus. Depending on the question at hand, one of several experimental preparations can be obtained from the sacculus. The simplest of these is the one-chamber preparation. Here the sacculus is immobilized in a chamber filled with artificial perilymph, a sodium-rich and high-calcium saline. This preparation enables the study of hair-cell currents and basic hair-bundle mechanics. A second configuration, the two-chamber preparation, can be used to study spontaneous hair-bundle movements. Here the apical side of hair cells is exposed to a potassium-rich and calcium-poor saline termed artificial endolymph, whereas the basolateral side is bathed in artificial perilymph. These two compartments mimic the *in vivo* arrangement of salines and provide an environment that allows hair bundles to oscillate spontaneously.

We describe in this paper the preparation of the bullfrog’s sacculus for biophysical study of its sensory hair cells. We first provide a detailed depiction of the isolation of this organ from the frog’s inner ear. We then describe both the one- and two-chamber experimental preparations and include representative results for each configuration.

## PROTOCOL

### 1. Pre-experimental preparation

#### 1.1) Solutions

1.1.1) Prepare the eugenol solution (2.5 g·L^−1^·kg^−1^ frog).
1.1.2) Prepare the saline solutions (Table 1).
  - For a one-chamber preparation, prepare artificial perilymph.
  - For a two chamber preparation, prepare both artificial perilymph and artificial endolymph.

### 2) Experimental tools

#### 2.1) Preparation of glass stimulation fiber

2.1.1) Narrow a borosilicate glass capillary with an electrode puller in a one-line pull with high heat and high velocity.
2.1.2) Load the pulled capillary into a solenoid-driven puller.
2.1.3) Bring the capillary’s tip toward a filament with a glass bead melted onto it until the capillary contacts the glass bead.
2.1.4) Turn on the filament to melt the tip of the capillary into the glass bead. Once a thin bridge of glass forms between the capillary and the bead, turn off the filament and around the same time activate the solenoid puller. This will pull the glass capillary at a right angle to its long axis and create a solid fiber.
2.1.5) Ensure that the fiber’s diameter is no more than 0.5-1 μm and that its length does not exceed 100-300 μm. If the fiber’s length exceeds this dimension, trim it with iris scissors. Use scissors that are already dull to avoid damaging your dissection tools.
2.1.6) To enhance optical contrast, sputter coat the fiber. Use a sputter coater with a gold-palladium source. Vertically load each fiber into a sputter coater with its tapered end towards the source.
2.1.7) Bring the fiber’s tip to a distance of 1-2 cm from the gold-palladium source.
2.1.8) Close the sputter coater’s chamber to form a seal and turn it on. Repeatedly flush out the ambient air with argon. After the air has been flushed, reduce the pressure within the chamber to 10 Pa (70 mTorr).
2.1.9) Sputter coat in 10-second pulses with 10-second delays over a course of 120 seconds. The fiber’s tip will darken over the duration of this protocol if the sputter coating was a success.

#### 2.2) Preparation of sharp microelectrodes

2.2.1) Pull a glass capillary with an internal filament using a one-line high-heat pull. Electrodes should have a resistance of 100-300 MΩ when filled with 3 M KCl.
2.2.2) Fill each electrode with 3 M KCI.
2.2.3) Bend each electrode’s tip with a microforge to render it perpendicular to the hair cell apical surface when mounted in the amplifier headstage^16^.

#### 2.3) Preparation of iontophoresis pipettes

2.3.1) Pull a glass capillary with an internal filament to a 50 MΩ tip.
2.3.2) Fill with concentrated solute (e.g. 500 mM gentamicin sulfate).

#### 2.4) Preparation of aluminum mounting square

2.4.1) Cut a 1 cm × 1 cm square of aluminum foil.
2.4.2) Perforate the foil in the center with the tip of a pithing rod or other sharp object.
2.4.3) Shape the perforation so that it is circular and approximately 1 mm in diameter.

#### 2.5) Preparation of vacuum grease-filled syringe

2.5.1) Remove the plunger from a 5 mL syringe.
2.5.2) Fill syringe from the back with vacuum grease.
2.5.3) Replace the plunger.

### 3. Extraction of inner-ear organs

3.1) Anesthetize an American bullfrog (Rana catesbeiana) by placing it in a small bucket containing eugenol anesthetic solution for 10 minutes. Adjust the volume of solution so that the frog’s humero-scapular joint lies just above the liquid-air interface.
3.2) Euthanize the anesthetized bullfrog by double pithing.
3.2.a. Grasp the anesthetized frog with one finger atop its nose and another below the jaw and rotate the frog’s head forward.
3.2.b. Rapidly plunge the pithing rod into the cranial vault through the foramen magnum, which is found at the midline between the frog’s occipital processes.
3.2.c. Slowly withdraw and rotate the rod until its tip departs from the foramen magnum. Force the rod caudally through the vertebral foramina to destroy the spinal cord.
3.2.d Confirm that the frog has been properly doubly pithed by observing that its lower extremities are extended.
3.3) Grasp the frog with a thumb atop the nose and first finger grasping the vomarine teeth for improved stability. Decapitate the frog by severing the temporomandibular joint bilaterally and subsequently cutting orthogonally to the rostrocaudal axis. To assure that the inner-ear organs within the skull remain intact, ensure that the cut is caudal to both tympana.
3.4) Using a stereodissection microscope, execute a midline cut through the palatal tissue from the vomarine teeth to the most posterior extent of the tissue (Figure 1A).
3.5) Sever and clear with horizontal cuts of the scalpel any muscle lying below the palatal tissue to reveal the posterior cartilage. With the muscle removed, the lollipop shape of the temporal cartilage forming the boundary of the otic capsule becomes visible.
3.6) Sever the columella at its point of contact with the otic capsule’s cartilage.
3.7) Repeatedly shave thin layers of this cartilage by making shallow horizontal cuts through it. Avoid deep cuts to prevent damage to the inner-ear organs. This opens the otic capsule found within the lollipop structure of the temporal cartilage (Figure 1B). Within the otic capsule are the frog’s inner-ear organs (Figure 1C).
3.8) Trim the posterior and lateral edges of the otic capsule, taking care not to damage the inner-ear organs. During the dissection, frequently flow saline over the inner ear organs to ensure that they remain submerged and hydrated.
3.9) Locate the two circular openings in the temporal bone at the medial connection of the cartilage to the midline. Cut downward through the most medial opening to rupture the temporal bone.
3.10) Make a second downward cut through the otic capsule at its postero-lateral edge. The piece of cartilage between this and the previous cut will be removed to provide access to the inner-ear organs.
3.11) Pry loose the cartilage between the cuts from steps 3.8 and 3.9 and cut it away from the otic capsule. This action severs the nearest semicircular canal, which can then be used as a handle for further manipulations.
3.12) Taking care not to touch the sacculus, sever the VIII^th^ cranial nerve.
3.13) Hold the ampulla of the nearest semicircular canal. Gently rotate the inner ear to expose the remaining two semicircular canals. Upon exposing each canal, sever it.
3.14) Holding the nerve or a semicircular canal ampulla, extract the inner ear from the head and place it in a dish filled with chilled oxygenated artificial perilymph. Removal of the inner ear permits visualization of the passages within the temporal bone through which the semicircular canals once passed (Figure 1D). *Repeat these steps to extract the second ear.*

**Figure 1:**
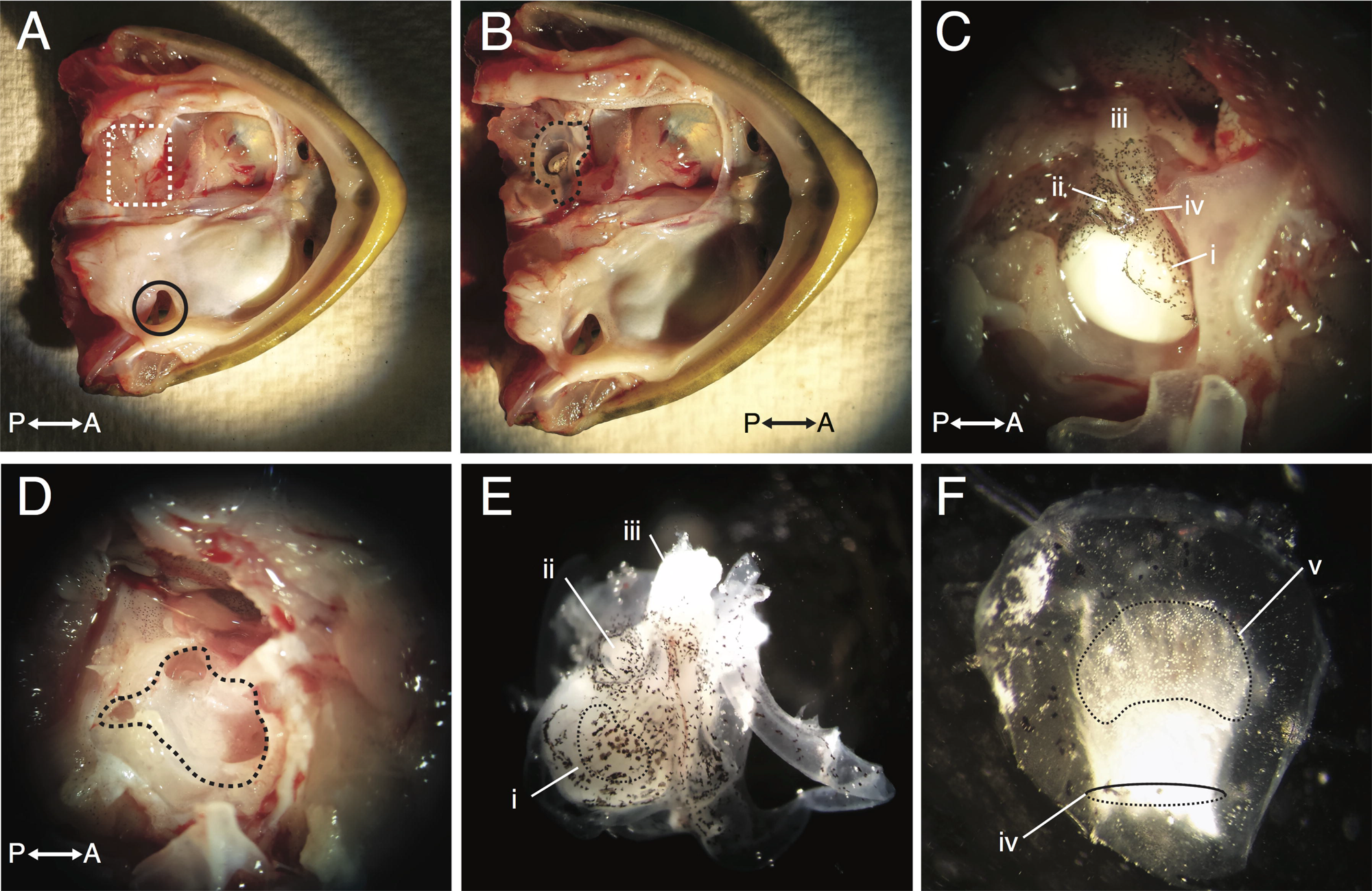
Dissection of the bullfrog’s inner ear. (A) Viewing the bullfrog’s upper palate from its ventral side permits identification of the Eustachian tube (black circle). Lateral reflection of the skin covering the right side of the upper palate reveals the location of the inner ear (dashed white box). (B) Removal of the cartilage on the ventral side of the frog’s temporal bone opens the otic capsule (dashed black line). (C) Displayed is a higher magnification image of the otic capsule, in which the sacculus, lagena, CN VIII, and saccular nerve can be readily identified. (D) A view of the otic capsule after removal of the inner-ear organs reveals the locations of the semicircular canals. (E) After removal of the isolated inner ear, the sacculus, lagena, and VIII^th^ cranial nerve (CN VIII) can readily be identified. (F) The isolated sacculus possesses a short stump of the saccular nerve and an otolithic membrane lying atop its sensory epithelium. Labels correspond to the (i) sacculus, (ii) lagena, (iii) CN VIII, (iv) saccular nerve, and (v) otolithic membrane. Axis labels P and A correspond respectively to the posterior and anterior directions.

### 4. One-chamber preparation

#### 4.1) Isolation of sacculus

4.1.1) Locate the sacculus by identifying its large white mass of otoconia and the saccular nerve that lies atop it (Figure 1E). Take care not to physically traumatize the sacculus during the following steps in order to maintain the integrity of saccular hair cells.
4.1.2) Trim the semicircular canals to render the inner ear more maneuverable.
4.1.3) Remove the perilymphatic cistern overlying the neural side of the sacculus. Make gentle cuts around the perimeter of the cistern. Next sever the small pillars of tissue that bridge the cistern’s membrane to the neural side of the sacculus.
4.1.4) Remove the lagena and its associated nerve.
4.1.5) Holding the saccular nerve, gently lift the sacculus and cut through the thin membrane of the otoconial sac. As otoconia spill out of the sac, free the sacculus by cutting around its perimeter.

> *After isolating the sacculus, the remaining inner-ear organs can be saved if desired. Removal of the sacculus eases identification of other structures, such as the amphibian and basilar papillae.*
4.1.6) Use scissors or forceps to carefully wipe away any otoconia remaining on the sacculus. Take care not to touch the sacculus during this process.
4.1.7) Trim any remaining otoconial-sac membrane from the sacculus’ edge. This membrane tends to adhere to plastic and glass surfaces and its removal minimizes challenges in handling the tissue.
4.1.8) Gently flow saline over the sacculus with a Pasteur pipette to remove any remaining otoconia (Figure 1F). *Repeat this procedure for the second sacculus, which is a mirror image of the first*.

#### 4.2) Digestion and mounting

4.2.1) Using the back end of a Pasteur pipette with its tip broken, transfer the isolated sacculi to a petri dish containing 3 mL of 67 mg·L^−1^ protease XXIV in artificial perilymph. This protease treatment digests the links tethering each kinociliary bulb to the otolithic membrane, enabling removal of the membrane without damaging the sensory hair bundles.
4.2.2) Incubate the tissue for 30 minutes at 22 °C (or 35 minutes at 21 °C).
4.2.3) Transfer each digested sacculus to an open-faced experimental chamber and secure the tissue with magnetic pins.
4.2.4) Carefully remove the otolithic membrane using a fine eyelash, taking care not to touch the saccular macula.

### 5. Two-chamber preparation

#### 5.1) Isolation of sacculus

Isolate the sacculus from the inner-ear organs as in Section 4.1.

#### 5.2) Mounting and digestion

5.2.1) Fill the mounting block with artificial perilymph and place the perforated aluminum foil on one opening, using two spots of vacuum grease to hold it in place and form a weak seal.
5.2.2) Transfer one sacculus to the foil and center it on top of the hole with the macula facing down and the nerve stump facing up.
5.2.3) Remove the saline surrounding the sacculus with a piece of twisted tissue (e.g. Kimwipe®). The saline should be wicked to dry the surface of the aluminum square surrounding the sacculus.
5.2.4) Use the Teflon applicator to apply cyanoacrylate glue to form a tight seal along the boundary between the edge of the sacculus and the aluminum square. Ensure that the entire circumference of the sacculus is covered with glue.

> *Proceeding too slowly allows the cyanoacrylate glue to creep over the neural side of the sacculus and eventually cover it. It is therefore imperative to complete this task rapidly.*
5.2.5) Place a drop of saline on top of the mounted tissue to cure the glue. A thin film of glue may form on top of the drop of saline; remove it with forceps.
5.2.6) Carefully remove the foil from the mounting block. Flip the mounted tissue over so that the macular side of the sacculus faces upward and float it on an artificial perilymph-filled petri dish.
5.2.7) Add a drop of protease XXIV solution on top of the macula and incubate for 30 minutes at 22 °C (or 35 minutes at 21 °C).
5.2.8) Fill the lower canal of the two-chamber apparatus with perilymph and place vacuum grease around the central chamber.
5.2.9) Place the foil-mounted sacculus on the lower chamber with its nerve facing the chamber’s surface. Add grease around the perimeter of the foil.
5.2.10) Place the upper chamber of the preparation on the foil, taking care to form a complete seal with the vacuum grease.
5.2.11) Fill the upper chamber with bubbled artificial endolymph and gently remove the otolithic membrane with an eyelash.

## REPRESENTATIVE RESULTS

The sensory epithelium of the bullfrog’s sacculus can be employed in various configurations to probe the physiology of hair cells. Because the tissue is relatively flat, it can be mounted in both one- and two-chamber preparations. The one-chamber configuration provides a simple setup for electrophysiological and micromechanical recordings of hair cells. The two-chamber preparation instead simulates both the endolymphatic and perilymphatic compartments on respectively the apical and basal sides of the sacculus. These compartments together provide a physiologically-relevant environment for the study of mechanotransduction by hair cells.

The sensitivity and transduction characteristics of hair cells underlie their electrical response to mechanical stimulation. To probe these features, we simultaneously recorded from an individual hair cell the position of its bundle and the cell’s receptor potential (Figure 2). We first attached a flexible glass fiber to the kinociliary bulb of a hair bundle to apply force pulses. We then measured the hair bundle’s displacement using a dual photodiode system^2^ (Figure 2A). We concurrently acquired the hair cell’s potential by impaling the cell with a sharp microelectrode. We obtained a displacement-response curve by plotting the peak voltage response elicited by each mechanical stimulus against the hair bundle’s corresponding displacement (Figure 2B). The hair cell’s electrical response saturates for both positive- and negative-displacement extrema. The reduction of membrane potential with negative displacement steps indicates the presence of a resting inward mechanotransduction current. This resting current is modulated by the action of Ca^2+^ on both fast and slow adaptation^17-21^.

**Figure 2:**
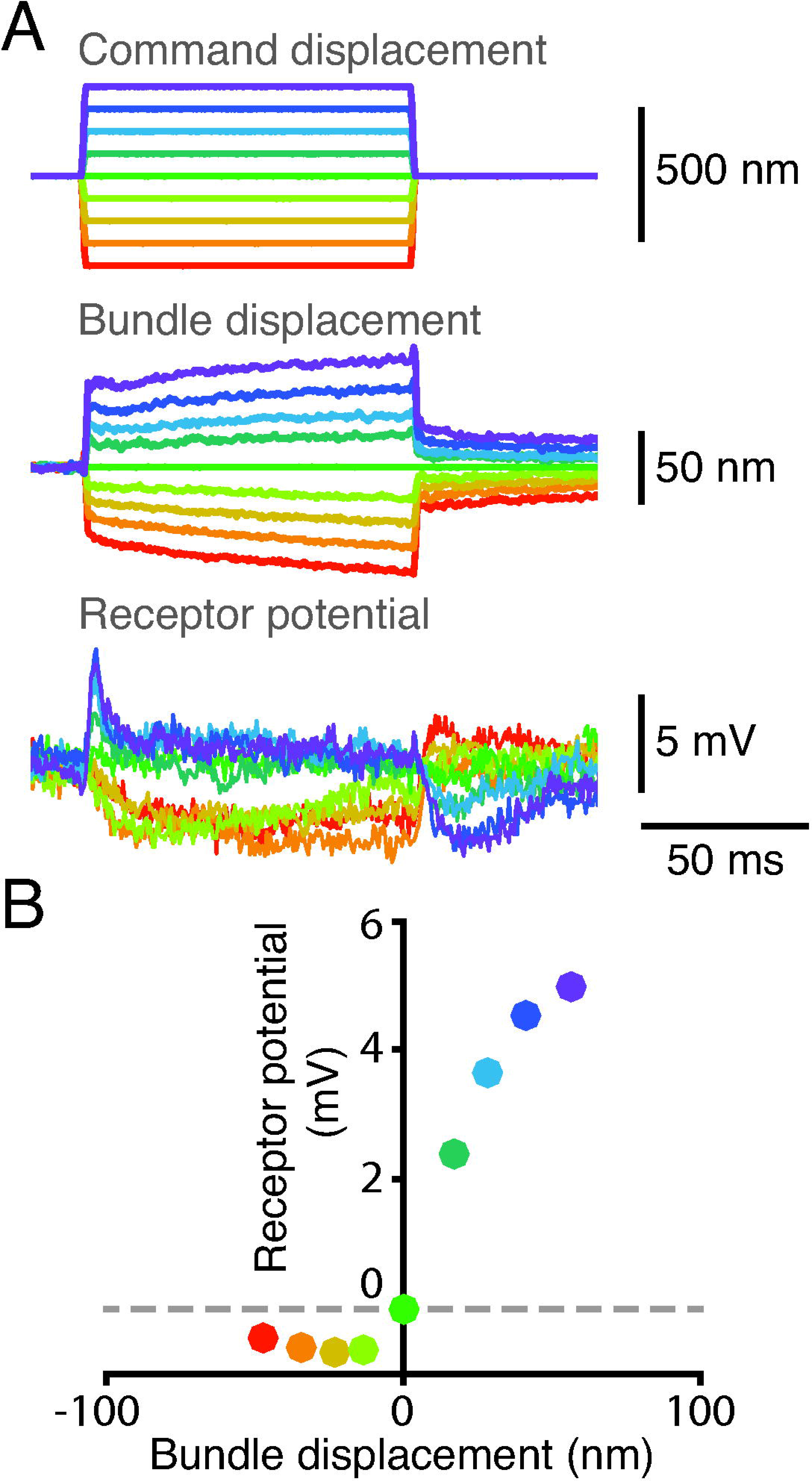
Displacement-response curve for a single hair cell. (A) The tip of a glass stimulus fiber was coupled to the kinociliary bulb of a hair bundle and the fiber’s base was subsequently displaced across nine discrete steps. The bundle’s position was tracked on a dual-photodiode system, and its receptor potential was simultaneously measured using a microelectrode whose output was passed through an amplifier in bridge mode. The electrode’s tip resistance was 95 MΩ and the bundle’s resting membrane potential was −47 mV. (B) A plot of the bundle’s receptor potential as a function of its displacement reveals a nonlinear relationship between the bundle’s response and its position. Each point corresponds to the mean potential and mean displacement over a 2.5 ms time window, beginning 2.5 ms after the onset of mechanical stimulation. Each color represents a set of time series corresponding to the same displacement pulse.

A hair cell’s behavior depends not only upon its electrical properties, but also upon the micromechanics of its sensory hair bundle. The two-chamber configuration mimics the separation of endolymph and perilymph *in vivo*, providing ideal conditions for the study of a hair bundle’s mechanics. Under these conditions and with the otolithic membrane removed, hair bundles can oscillate spontaneously^6^. Here we employed the two-chamber preparation to assess the micromechanics of individual bundles. We recorded the spontaneous oscillations of a hair bundle by casting its shadow on a dual-photodiode displacement monitor (Figure 3). To assess the role of the aminoglycoside antibiotic gentamicin on mechanotransduction, we iontophoretically released gentamicin directly onto the hair bundle (Figure 3). The concentration of released gentamicin rises proportionally with the current passed through the micropipette. Gentamicin inhibits a hair bundle’s oscillations and induces a static offset of the bundle towards its tall side. These effects reflect the role of gentamicin as an open-channel blocker that maintains the open state of mechanotransduction channels while blocking their permeation pore. Iontophoresis of charged chemicals permits localized and quantifiable release of chemicals at various concentrations in the absence of fluid flow-induced mechanical disruption and is thus ideally suited for the study of mechanosensitive organelles such as the hair bundle^22^.

**Figure 3:**
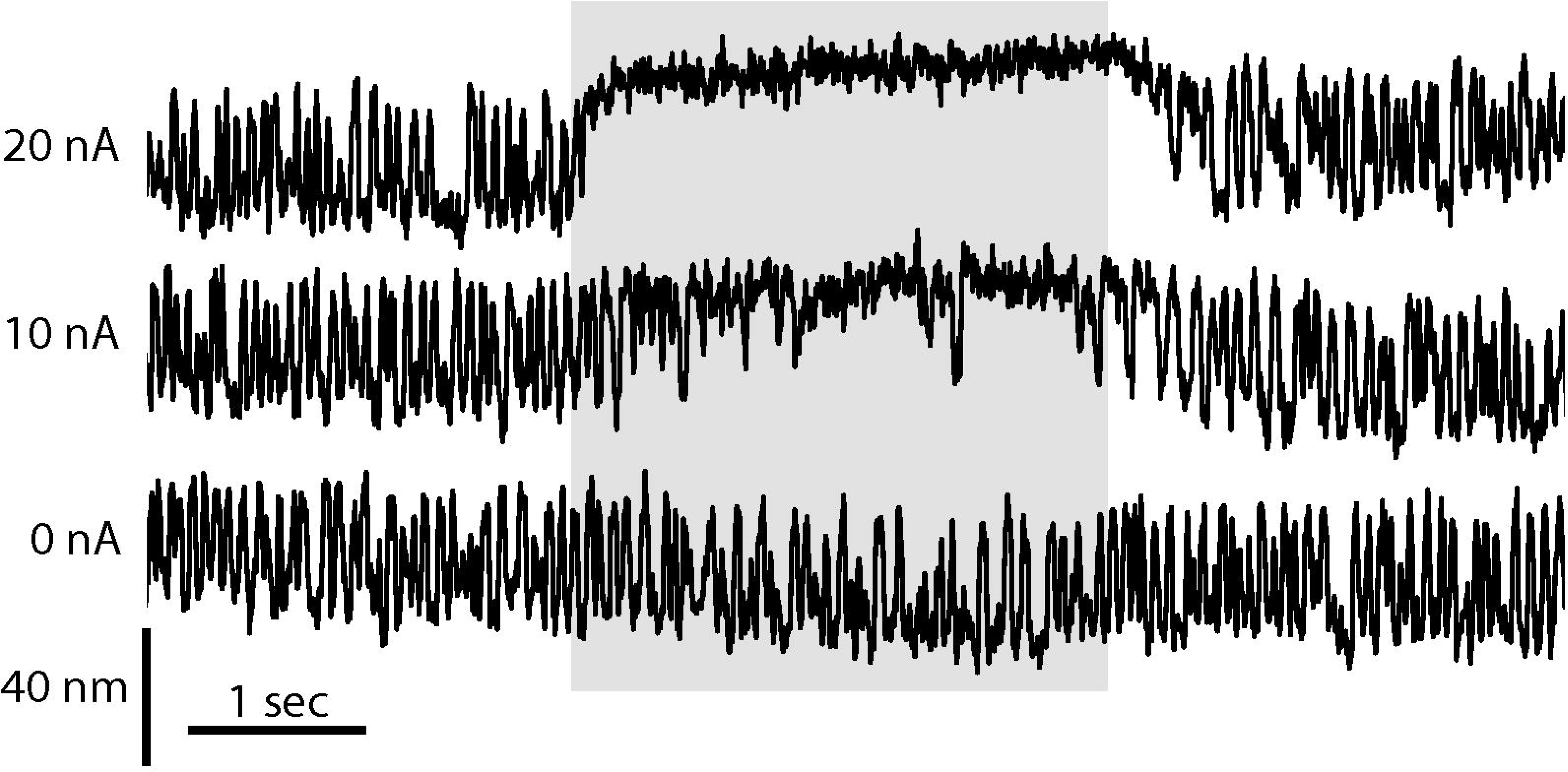
Effect of gentamicin on spontaneous hair-bundle oscillation. The spontaneous motion of a hair bundle in a two-chamber preparation was recorded using a dual photodiode system. In the absence of iontophoretic release of gentamicin (0 nA), the hair bundle displays symmetric oscillations. As the magnitude of current passed through an iontophoretic pipette filled with 500 mM gentamicin sulfate grows (10 nA, 20 nA), the frequency of hair-bundle excursions falls in a dose-dependent manner and the bundle is offset towards its tall edge for longer periods of time.

A hair bundle’s spontaneous motion arises from the interplay between adaptation and a nonlinearity^7,8,23,24^. This nonlinearity corresponds to nonlinear bundle stiffness. Hair bundles have been shown to exhibit nonlinear stiffness in the vestibular^2^, auditory^25^, and lateral-line systems^26^. A hair bundle’s active process depends on the nature of this nonlinearity, and measurement of the bundle’s stiffness reveals additional mechanisms underlying the dynamics of the active process.

We directly measured the stiffness of an individual hair bundle from the bullfrog’s sacculus (Figure 4). To achieve this, we coupled the tip of a flexible glass fiber to the hair bundle’s kinociliary bulb (Figure 4A). We delivered forces to the hair bundle by displacing the fiber’s base. The force exerted onto the hair bundle by the stimulus fiber corresponds to the difference between the displacements of the fiber’s base and tip, multiplied by the fiber’s stiffness^2,27^. Delivering pulses across a range of forces reveals a relationship between the force exerted onto the bundle and the bundle’s ensuing displacement. The slope of this force-displacement relation corresponds to the hair bundle’s stiffness (Figure 4A).

**Figure 4.**
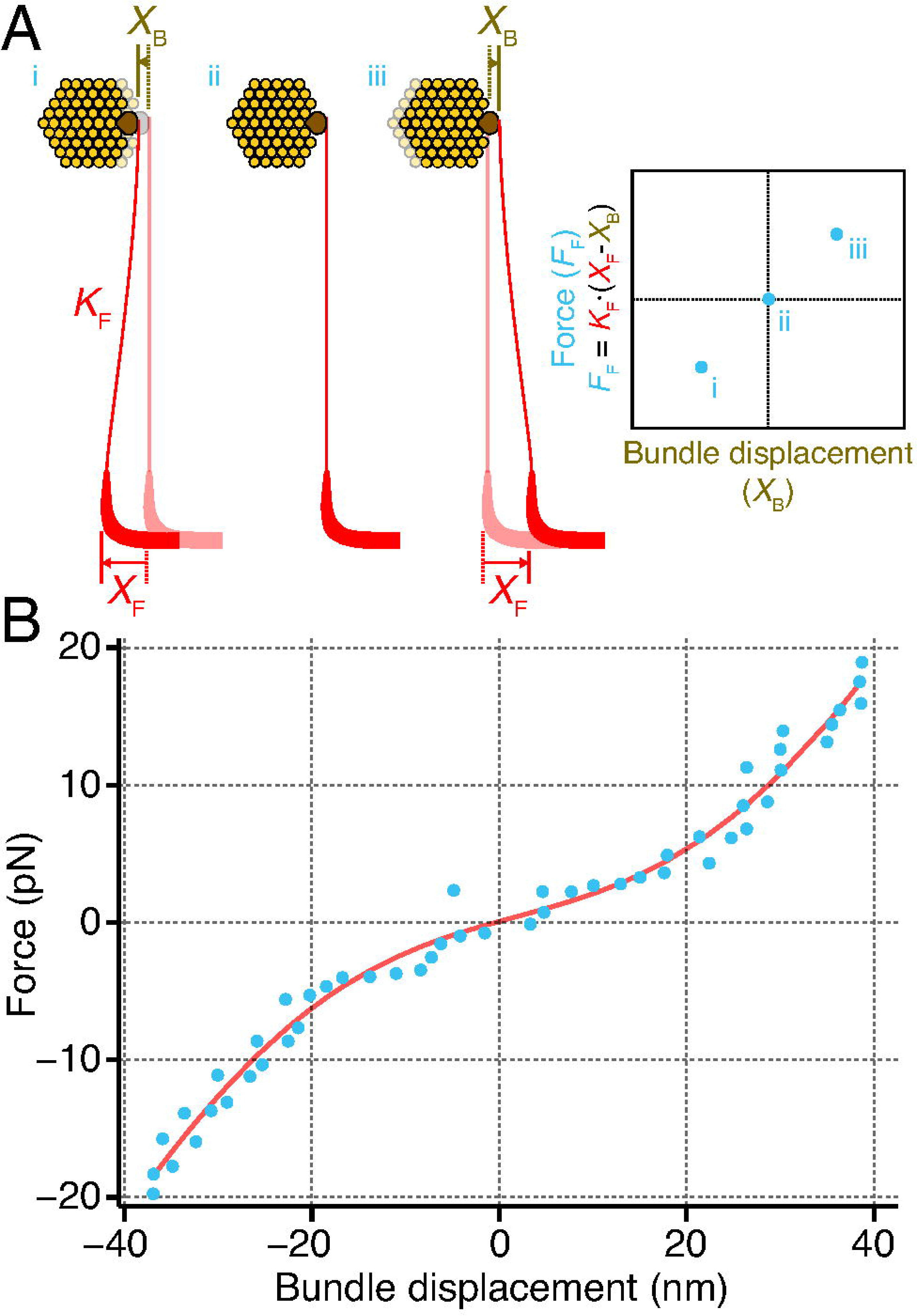
Calculating a hair bundle’s stiffness. (A) A stimulus fiber (red) of stiffness K_F_ is coupled to the kinociliary bulb (brown) of an individual hair bundle (yellow). Displacing the base of the fiber a known distance X_F_ causes the bundle to move a distance X_B_. The difference between the displacements of the fiber and the bundle is proportional to the force exerted onto the bundle by the stimulus fiber, *F*_F_. Repeating this across a range of forces yields a force-displacement relation (right), the slope of which corresponds to the hair bundle’s stiffness. (B) An individual bundle was subjected to force pulses of increasing magnitude and its displacement within the first 50 ms after the pulse onset was measured (blue points). Here the hair bundle displays a nonlinear stiffness over a range of approximately 20 nm around its resting position. The red curve corresponds to a fit to the relation *F* = *k*\**X* − 60\**z*\*(1/(1+exp(-*z*\*(*X*−*X*_0_)/(*k*_B_\**T*))) + *F*_0_, in which *F* is the force applied to the bundle, *X* is the bundle’s displacement, *k* = 790 ± 51 μN·m^−1^ is the bundle’s linear stiffness when all channels are either closed or open, *z* = 0.43 ± 0.04 pN is the force of a single gating spring, *X*_0_ = 2 ± 1.9 nm is the bundle’s position at which 50% of its channels are open, *k*_B_ is Boltzmann’s constant, T is the temperature, and *F*_0_ = 11.7 ± 1.3 pN is an offset force. The fit possesses a coefficient of determination of 0.98. The stimulus fiber had a stiffness of 107 μN·m^−1^.

This method allowed us to measure an individual bundle’s stiffness as a function of its deflection (Figure 4B). The force-displacement curve displays a nonlinear relationship, revealing a nonlinear stiffness of the bundle over a range of about 20 nm around its resting position. Outside this range, the hair bundle behaves like a Hookean material, its stiffness is linear for large-magnitude deflections.

These results demonstrate the versatility of the bullfrog sacculus in the study of hair-cell physiology. Using these and other preparations, one can explore mechanotransduction at multiple stages in the transmission of information from the bundle toward the brain.

## DISCUSSION

Within the bullfrog’s sacculus lie several thousand easily-accessible sensory hair cells. Here we demonstrate extraction and preparation of the sacculus for one- and two-chamber recordings. These two preparations permit both micromechanical and electrophysiological studies of hair cells and their associated bundles. Because the tissue can survive for several hours with frequent replacement of oxygenated saline, experiments may continue for long durations. Hair cells in these preparations typically remain viable for microelectrode recording for up to 6 hours after dissection, while hair bundles oscillate spontaneously for up to 24 hours after extraction.

Successful extraction and mounting of the sacculus hinges upon surmounting several common challenges. First, direct contact with the apical surface of the saccular macula should be avoided throughout the preparation procedure. The saccular nerve provides a convenient handle for safe manipulation of the sacculus. Once freed from the remainder of the inner-ear organs, the sacculus should be transferred using a large-bore pipette while remaining immersed in fluid to avoid mechanical damage to its sensory epithelium. The removal of otoconia from the macular surface must be completed without mechanical damage to hair cells. Because the otoconia lie directly atop the macula, hair cells can be damaged by accidental contact between dissection tools and the otolithic membrane while removing otoconia. To avoid damage, we recommend that the gelatinous mass of otoconia be held at a location far from the macula and removed as a single mass. This avoids fragmentation of the otoconial mass into numerous clusters, each of which would be individually extracted. If small clusters of otoconia remain they can be removed with gentle fluid pressure delivered by a Pasteur pipette. A final challenge involves the formation of a tight seal between the sacculus and aluminum mounting square in the two-chamber preparation. Employing a square with a perforation small enough to allow overlap of about 100 μm between the sacculus and the surrounding aluminum permits complete sealing of the tissue. The glue should be brought into contact with approximately 100 μm of saccular tissue around its perimeter to form a tight seal.

The concentration of free Ca^2+^ is an important consideration in the study of hair cells. Ca^2+^ regulates both fast and slow adaptation, thus determining the kinetics of the mechanotransduction apparatus and the characteristics of the hair bundle’s active-process phenomena, including spontaneous bundle motion^8,23^. Endolymphatic calcium *in vivo* is present at 250 μM, therefore the most physiologically relevant kinetics are assessed at this concentration (Maunsell J. H. R., R. Jacobs, and A. J. Hudspeth. Unpublished observations^16^). However, microelectrode recordings from hair cells require an external calcium concentration exceeding 2 mM for proper sealing of the cellular membrane around the microelectrode. It is therefore imperative to use a high-calcium saline for these experiments. Finally, one may wish to study the effects of external calcium upon mechanotransduction using a variety of calcium concentrations. In these cases it is important to remember that calcium concentrations below 1 μM typically lead to tip-link rupture and irreversible loss of transduction^28^.

The two experimental preparations described here allow for a range of biophysical measurements on hair cells. However, additional measurements can be made with slight modifications to these preparations. In the folded saccular preparation, hair bundles are visualized laterally. Imaging hair-bundle motion from this vantage point reveals coherent motion of both short and tall stereocilia^29^. Here the saccular macula is first separated from its underlying tissue and subsequently folded along the axis defined by the saccular nerve such that hair bundles face outwards and are visualized laterally at the crease. A second modification, hair-cell dissociation, enables the study of both the hair cell’s bundle and its soma. Hair cells are mechanically dissociated onto a glass slide for imaging and electrophysiological recording^30^. Finally, hair cells can be extruded from the epithelium by following a similar dissociation protocol but without the mechanical dissociation step. This treatment results in hair cells that automatically extrude out of the epithelium, providing basolateral access for electrophysiological recordings while minimizing mechanical damage. These preparations and their many modifications demonstrate the versatility of the frog sacculus as a model system for the biophysical study of sensory hair cells.

## Supporting information

Materials List

## ACKNOWLEDGMENTS

The authors wish to acknowledge Dr. A. J. Hudspeth for funding and expertise in developing the preparations described in this paper. We also wish to thank Brian Fabella for creating and maintaining much of the custom equipment and software used in this protocol.

J. B. A. is supported by grant F30DC014215, J. D. S. is supported by grant F30DC013468, and both J. B. A. and J. D. S. are supported by grant T32GM07739 from the National Institutes of Health.

---

**Table 1. Solutions for dissection and experimental preparation**. Displayed in this table are the recipes for artificial perilymph and artificial endolymph solutions used in dissection and in one- or two-chamber preparations. The solutions should be brought to pH 7.2-7.4 with about 2 mL of NaOH (perilymph) or 2 mL KOH (endolymph). The osmotic strength should read approximately 230 mmol·kg^−1^ due to incomplete ionic dissociation.

