## Supplementary material for "Physiological preparation of hair cells from the sacculus of the American bullfrog (*Rana catesbeiana*)": Materials List

Materials List for:

URL: http://www.jove.com/video/55380
DOI: doi:10.3791/55380

### Materials

| Name | Company | Catalog Number | Comments |
|---|---|---|---|
| **Common to both preparations** | | | |
| Stereo-dissection microscope | Leica | MZ6 | Other sources can be used |
| Tricaine methanesulfonate | Sigma | E10521 | Other sources can be used |
| Metal pithing rod | Fine Science Tools | 10140-01 | |
| Vannas spring scissors | Fine Science Tools | 15000-03 | |
| Dumont #5 forceps | Fine Science Tools | 11252-20 | |
| Glass Pasteur pipette and bulb (x2) | Fisher Scientific | 22-042816 | |
| Fine eyelash mounted on a hypodermic needle | Fisher Scientific | 22-557-172 | |
| Dow-corning vacuum grease | Fisher Scientific | 14-635-5C | |
| Syringe for vacuum grease | Fisher Scientific | 14-829-45 | Other sources can be used |
| 35 mm Petri dish (x2 - 3) | Fisher Scientific | 08-772A | Other sources can be used |
| Micropipette puller | Sutter | P-97 or P-2000 | |
| 120 V Solenoid puller | | | Home-made, see parts list |
| Sputter coater | Anatech USA | Hummer 6.2 | |
| Current source for iontophoresis | Axon Instruments | AxoClamp 2B | Other sources can be used |
| Piezoelectric actuator | Piezosystem Jena | P-150-00 | |
| Amplifier for piezoelectric actuator | Piezosystem Jena | ENV800 | |
| Borosilicate glass capillary | World Precision Instruments | 1B120F-3 | |
| **Name** | **Company** | **Catalog Number** | **Comments** |
| **For one-chamber preparation** | | | |
| Microelectrode amplifier | Axon Instruments | AxoClamp 2B | Can be used for iontophoresis and microelectrode recordings simultaneously |
| Magnetic pins (x2) | | | Home-made, see parts list |
| Open-top chamber with magnetic sheet | | | Home-made, see parts list |
| **Name** | **Company** | **Catalog Number** | **Comments** |
| **For two-chamber preparation** | | | |
| Upper chamber | | | Supplementary file 1 |
| Troughed lower chamber | | | Supplementary file 2 |
| Aluminum foil | Fisher Scientific | 01-213-100 | Other sources can be used |
| Mounting block | | | Supplementary file 3 |
| Wooden applicator sticks | Fisher Scientific | 23-400-112 | Other sources can be used |
| Teflon sheet | McMaster-Carr | 8545K12 | For teflon applicator |

Copyright © 2017 Creative Commons Attribution-NonCommercial-NoDerivs 3.0 Unported License

| | | | |
|---|---|---|---|
| Cyanoacrylate glue | 3M | 1469SB | |
| Lab tissues (Kimwipes) | Fisher Scientific | 06-666A | Other sources can be used |
| Gentamicin sulfate | Sigma-Aldrich | G1914 | Other sources can be used |
| Quick-setting epoxy | McMaster-Carr | 7605A18 | |
| 18 mm glass coverslips | Fisher Scientific | 12-546 | Other sources can be used |
| **Name** | **Company** | **Catalog Number** | **Comments** |
| **Saline components** | | | |
| NaCl | Fisher Scientific | S271-3 | Other sources can be used |
| KCl | Sigma-Aldrich | P4504-500G | Other sources can be used |
| $CaCl_2 \cdot 2H_2O$ | Fisher Scientific | 10035-04-8 | Other sources can be used |
| HEPES | Sigma-Aldrich | H3375-100G | Other sources can be used |
| D-(+)-glucose | Sigma-Aldrich | G7021 | Other sources can be used |
| **Name** | **Company** | **Catalog Number** | **Comments/Description** |
| **Parts lists for home-made equipment** | | | |
| **Solenoid puller** | | | |
| Solenoid | Guardian Electric | A420-065426-00 | Other sources can be used |
| Foot-pedal switch | Linemaster | T-51-SC36 | Other sources can be used |
| Pipette holder | World Precision Instruments | MEH900R | Other sources can be used |
| Coarse manipulator | Narishige Group | MM-3 | Other sources can be used |
| Platinum wire | Alfa Aesar | 25093 | Other sources can be used |
| Power supply | Leica | Z050-261 | Other sources can be used |
| **Name** | **Company** | **Catalog Number** | **Comments/Description** |
| **Magnetic pins** | | | |
| Epoxy | McMaster-Carr | 7556A33 | Other sources can be used |
| 1 mm thickness aluminum | McMaster-Carr | 89015K45 | Other sources can be used |
| Insect pins | Fine Science Tools | 26000-40 | Other sources can be used |
| **Name** | **Company** | **Catalog Number** | **Comments/Description** |
| **Open-top magnetic chamber** | | | |
| Flexible magnetic strip | McMaster-Carr | 5759K75 | Other sources can be used |
| 1 mm thickness aluminum | McMaster-Carr | 89015K45 | Other sources can be used |

Copyright © 2017 Creative Commons Attribution-NonCommercial-NoDerivs 3.0 Unported License
